# TOW links TIR1/AFB-mediated signalling with Receptor-Like Kinases in auxin canalization

**DOI:** 10.1101/2025.04.25.650570

**Authors:** Mingyue Li, Nikola Rydza, Ewa Mazur, Gergely Molnár, Tomasz Nodzyński, Jiří Friml

## Abstract

Auxin canalization is a self-organizing process that governs the flexible formation of vasculature by reinforcing the formation of auxin transport channels. A key prerequisite is the feedback between auxin signaling and directional auxin transport, mediated by PIN transporters. Despite the developmental importance of canalization, the molecular components linking auxin perception to the regulation of PIN auxin transporters remain poorly understood. Here, we identify TOW, a novel and essential component of auxin canalization that links intracellular auxin signaling with cell surface auxin perception. TOW is regulated downstream of TIR1/AFB–Aux/IAA–WRKY23 transcriptional auxin signaling. *tow* mutants exhibit defects in regeneration and *de novo* vasculature formation, along with impaired formation of polarized, PIN-expressing auxin channels. At the subcellular level, these mutants display disrupted auxin-induced PIN polarization and altered PIN endocytic trafficking dynamics. TOW localizes to the Golgi, trans-Golgi network, and predominantly to the plasma membrane, where it interacts with receptor-like kinases involved in auxin canalization, including the TMK1 auxin co-receptor and the CAMEL–CANAR complex.

Together, our findings identify TOW as a molecular link between intracellular and cell surface auxin signaling mechanisms that converge on PIN trafficking and polarity, providing new insights into how auxin signaling regulates directional auxin transport for the self-organizing formation of vasculature during flexible plant development.

## Introduction

Plants have evolved delicate mechanisms to adapt their architecture to environmental conditions. Much of this adaptive development is achieved through self-organization of patterning processes, including the integration of new organs into the pre-existing vascular network, the emergence of complex leaf venation patterns, and the flexible regeneration of vasculature around wounds. Previous studies have identified auxin as the primary phytohormone involved in these processes, in particular, in a spontaneous formation and differentiation of vascular strands in different contexts (Sachs, 1975). The auxin canalization hypothesis was proposed, suggesting that the directional movement of an auxin-dependent signal through a field of cells enhances the ability of individual cells to transport auxin. This positive feedback mechanism linking auxin and its transport leads to the gradual formation of narrow auxin transport channels (Hajný et al., 2022). While the canalization hypothesis is both mechanistically fascinating and developmentally crucial - and is well supported by mathematical modeling (Bennett et al., 2014; Hajný et al., 2022) - the underlying genetic basis remains largely unknown.

Polarly localized PIN auxin efflux transporters are key components that facilitate directional cell-to-cell auxin flow in plants (Luschnig and Friml, 2024). PIN polarity, which is key determinant of auxin flow directionality (Wiśniewska et al., 2006), is extensively regulated by phosphorylation and depends of the endosomal trafficking (Glanc et al., 2018; Glanc et al., 2021); both processes being targeted by many endogenous and environmental signals (Luschnig and Friml, 2024). It has been repeatedly demonstrated that PIN polarity can be dynamically rearranged also by auxin (Sauer et al., 2006; Balla et al., 2011; Prát et al., 2018). Auxin treatment promotes the relocation of PIN proteins from the basal to the lateral side of root cells in a tissue-specific manner. Intriguingly, mutants with defects in this auxin-induced PIN lateralization exhibit typically abnormal leaf venation and vasculature formation (Mazur et al., 2016; Tejos et al., 2018; Hajný et al., 2020), reinforcing the idea that auxin feedback on PIN polarity is essential for vascular patterning, as proposed by the auxin canalization hypothesis. Yet, the molecular players coordinating PIN polarity, auxin transport, and endosomal trafficking during canalization remain largely unknown.

The canonical auxin signaling mechanism has been well characterized, involving the TIR1/AFB (TRANSPORT INHIBITOR RESPONSE 1/AUXIN-SIGNALING F-BOX), Aux/IAA (AUXIN/INDOLE-3-ACETIC ACID), and ARF (AUXIN RESPONSE FACTOR) components (Vanneste et al. *in press*). Mutants with defects in auxin signaling typically display disrupted vascular strand formation and abnormal leaf vein patterns (Sauer et al., 2006; Prát et al., 2018; Mazur et al., 2020). Transcriptional profiling of SCF^TIR1^-Aux/IAA-ARF signaling identified WRKY DNA-BINDING PROTEIN 23 (WRKY23) as a downstream regulator required for PIN lateralization and auxin canalization (Prát et al., 2018). These findings suggest that the SCF^TIR1^-Aux/IAA-WRKY23 signaling pathway is essential for auxin-mediated PIN polarity regulation during canalization. However, WRKY23 as a transcriptional regulator is not directly involved in trafficking or polarization processes, raising a question of identifying direct molecular players that mediate coordinated PIN polarity rearrangements during auxin canalization and beyond.

Among the downstream components of SCF^TIR1^-Aux/IAA-WRKY23 involved in PIN polarity regulation, the leucine-rich repeat receptor-like kinase (LRR-RLK), named CANALIZATION-RELATED AUXIN-REGULATED MALECTIN-TYPE RLK (CAMEL) has been identified (Hajný et al., 2020). Together with its interactor, CANALIZATION-RELATED RLK (CANAR), CAMEL phosphorylates PIN proteins, leading to their repolarization.

Notably, besides intracellular TIR1/AFB-mediated auxin signaling, also cell surface auxin perception mediated by AUXIN BINDING PROTEIN1 (ABP1) and its LRR-RLK interactors TRANSMEMBRANE KINASEs (TMKs) is essential for auxin canalization-mediated processes such as *de novo* vasculature formation and regeneration (Friml et al., 2022). The ABP1-TMK signaling-dependent regulation of auxin transport may occur via direct interaction of TMKs with PIN auxin transporters and their subsequent phosphorylation (Rodriguez et al., 2022; Wang et al., 2022). Nonetheless, how ABP1-TMK and TIR1/AFB auxin perception mechanisms are integrated with each other and with CAMEL-CANAR-mediated canalization mechanisms remain unclear.

In this study, we identified TOW, a previously uncharacterized membrane protein essential for auxin canalization, PIN polarity and trafficking. TOW is a downstream effector of the SCF^TIR1^–Aux/IAA–WRKY23 signaling and interacts with both, ABP1-TMK1 and CAMEL/CANAR signaling modules. Collectively, our findings establish TOW as a pivotal integrator of intracellular and cell surface auxin signaling with RLK-mediated PIN trafficking and polarity control, which is essential for executing auxin canalization during vascular development.

## Results

### TOW Is a Downstream Target of SCF^TIR1^-Aux/IAA-WRKY23 Signaling

To identify additional molecular factors involved in auxin canalization-dependent, self-organizing vasculature formation downstream of SCF^TIR1^-Aux/IAA-WRKY23, we analyzed transcriptomics data of auxin-responsive genes (Okushima et al., 2005), Aux/IAA-dependent genes (Prát et al., 2018), and WRKY23-dependent genes (Hajný et al., 2020) from previously published studies. The overlap between the auxin-responsive dataset and the Aux/IAA-induced gene set revealed a group of 245 genes that are auxin-induced and dependent on SCF^TIR1^-Aux/IAA-ARF signaling. Further comparison with WRKY23-dependent microarray data led to a final list of 53 genes as potential targets of the SCF^TIR1^-Aux/IAA-WRKY23 signaling (Supplemental Figure 1A). This group includes the AGCIII kinase *PINOID (PID;* Christensen et al., 2000), a well-characterized regulator of PIN polarity, along with numerous well-known components of transcriptional auxin signaling including *AuxIAAs*, auxin response factors (*ARFs; Okushima et al., 2005)*, and various auxin-responsive genes such as *SAUR*s, *LBD*s, and auxin biosynthetic enzymes like *GH3.1, GH3.3*, and *DFL1*. After filtering out these known components, we identified the uncharacterized gene *AT3G09280* as a potential novel factor required for auxin canalization (Supplemental Figure 1A). We named this gene *Target Of WRKY23 (TOW)*. Quantitative RT-qPCR confirmed that *TOW* transcription is significantly upregulated by IAA in a time- and dose-dependent manner; showing even faster and more sensitive response than *WRKY23* (Supplemental Figure 1B and 1C). This upregulation is largely compromised in the *arf7 arf19* double mutant, confirming that TOW is a downstream target of SCF^TIR1^-Aux/IAA-ARF signaling (Figure 1A). TOW transcription is also increased when the WRKY23 is conditionally activated in dexamethasone (DEX)-inducible line (Figure 1B).

**Figure 1.**
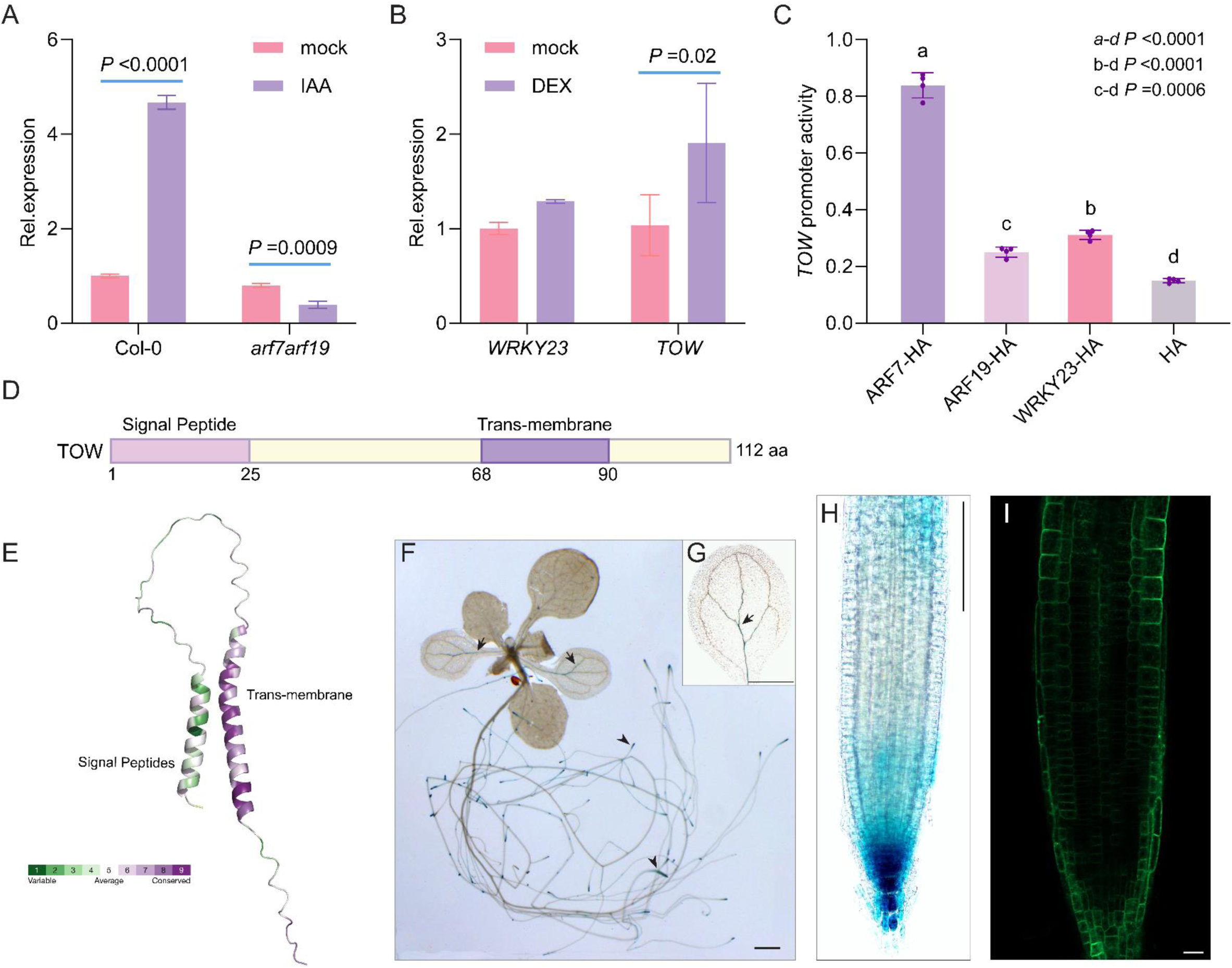
Auxin-inducible membrane protein TOW transcriptionally regulated by ARF and WRKY23 factors. **(A, B)** Quantitative RT-PCR analysis showing IAA-induced, ARF-dependent (A) as well as WRK23-dependent (B) regulation of *TOW* transcription. For all qRT-PCR experiments, gene expression levels were normalized to *PP2A*. *P* values were calculated using two-way ANOVA followed by Tukey’s post hoc test. Data are representative of three biological replicates and presented as mean ± SD. **(C)** Transactivation of *TOW* promoter by ARF and WRK23 after treatment with 1µM IAA for 24 hr in *Nicotiana benthaminana.* Data are mean ± SD, n = 3 independent biological replicates. *P* values were determined using one-way ANOVA with Tukey’s post hoc test. **(D)** Predicted domain structure of the TOW protein. **(E)** Evolutionary conservation scores of individual amino acids projected onto the predicted 3D structure of TOW. **(F-H)** Histochemical visualization of GUS activity (blue) in *TOW::GUS* reporter lines. Twelve-day-old seedling stained with GUS substrate for 6 hr (n = 8). Arrow, cotyledon leaf veins; arrowheads, root tips (F). Cotyledon GUS staining (n = 8) (G). Tip of primary root (n = 8) (H). **(I)** Subcellular localization of *pTOW::TOW-GFP* in the primary root tip (n = 10). Scale bars: 1 mm (F and G), 100 µm (H), 10 µm (I).

Consistent with the induction of TOW transcription by ARFs and WRKY23, three auxin-responsive elements (AuxREs) and three W-box binding motifs are annotated in the TOW promoter region, according to the JASPAR database (Khan et al., 2018) (Supplemental Figure 1D). Moreover, transactivation of *TOW* promoter by ARF7, ARF19, and WRKY23, respectively, after treatment with 1 µM IAA in *Nicotiana benthamiana* (Figure 1C), further confirmed that TOW is a direct downstream target of SCF^TIR1^-Aux/IAA-ARF and WRKY23 signaling.

### *TOW* Encodes a Small Transmembrane Protein Expressed in Developing Vasculature

Phylogenetic analysis based on the TOW amino acid sequence revealed that TOW is a plant-specific protein present exclusively in angiosperms (Supplemental Figure 2). In *Arabidopsis thaliana*, it is encoded by a single-copy gene. The most ancient identifiable ortholog is found in *Amborella trichopoda*, the earliest-diverging extant lineage of flowering plants (Supplemental Figure 2). The restriction of TOW to flowering plants suggests its involvement in processes specific to angiosperm development and physiology, possibly reflecting an evolutionary innovation linked to the emergence of complex vascular tissues or specialized auxin transport mechanisms.

Analysis of the TOW amino acid sequence revealed a predicted signal peptide at the N-terminus and a putative transmembrane domain near the C-terminus (Figure 1D). Evolutionary rate analysis showed high conservation scores within the transmembrane domain (Figure 1E), suggesting its functional importance. To investigate subcellular localization, we generated transgenic Arabidopsis plants expressing a *pTOW::TOW-GFP* construct. TOW-GFP predominantly localized to the plasma membrane in root epidermal and columella cells (Figure 1I), supporting its identity as a transmembrane protein.

To examine TOW expression patterns, we generated transgenic lines expressing a *GUS* (*β-glucuronidase*) reporter driven by a 2 kb *TOW* promoter. GUS staining revealed *TOW* expression in cotyledon leaf veins, petioles, and the root tips of both primary and lateral roots (Figure 1F–H). This expression pattern overlaps with, and partially precedes the formation of differentiating vascular strands and coincides with PIN1-expressing vascular tissues, consistent with a role for TOW in auxin-induced, PIN1-mediated vascular development.

### TOW Mediates Leaf Venation and the Underlying PIN1 Polarized Channel Formation

Auxin canalization underlies several developmental processes, including leaf vein formation (Scarpella et al., 2006), vascular regeneration after wounding (Mazur et al., 2016), and apical dominance (Balla et al., 2016). To investigate whether TOW is required for these processes, we examined the leaf venation pattern in cotyledons of Col-0 and *tow* mutants. Four independent *tow* alleles were generated using the CRISPR/Cas9 system (Supplemental Figure 3A). Among them, *tow-C1* and *tow-C3* carry in-frame deletions, while *tow-C2* and *tow-C4* harbor frameshift mutations that result in premature stop codons (Supplemental Figure 3A).

While the majority of cotyledons in Col-0 displayed regular, four-loop venation, approximately 20.6% showed mild irregularities (Figure 2A and 2B). In contrast, *tow* mutants exhibited a significant increase in defective vein patterns, with 34.3%–40.7% of cotyledons affected. Notably, *tow-C1* and *tow-C3* displayed reduced vein loop numbers (21.2% and 16.7% of cotyledons), whereas *tow-C2* and *tow-C4* showed a higher frequency of disconnected bottom loops (*tow-C2*, 20.1%; *tow-C4*, 13.3%) (Figure 2B, Supplemental Figure 3B). These defects were largely rescued by introducing a *pTOW::TOW-GFP* construct into the *tow* background, restoring regular venation in 76.8% of cotyledons—comparable to Col-0 levels (Figure 2B). These results show that the TOW function is essential for proper leaf vein formation.

**Figure 2.**
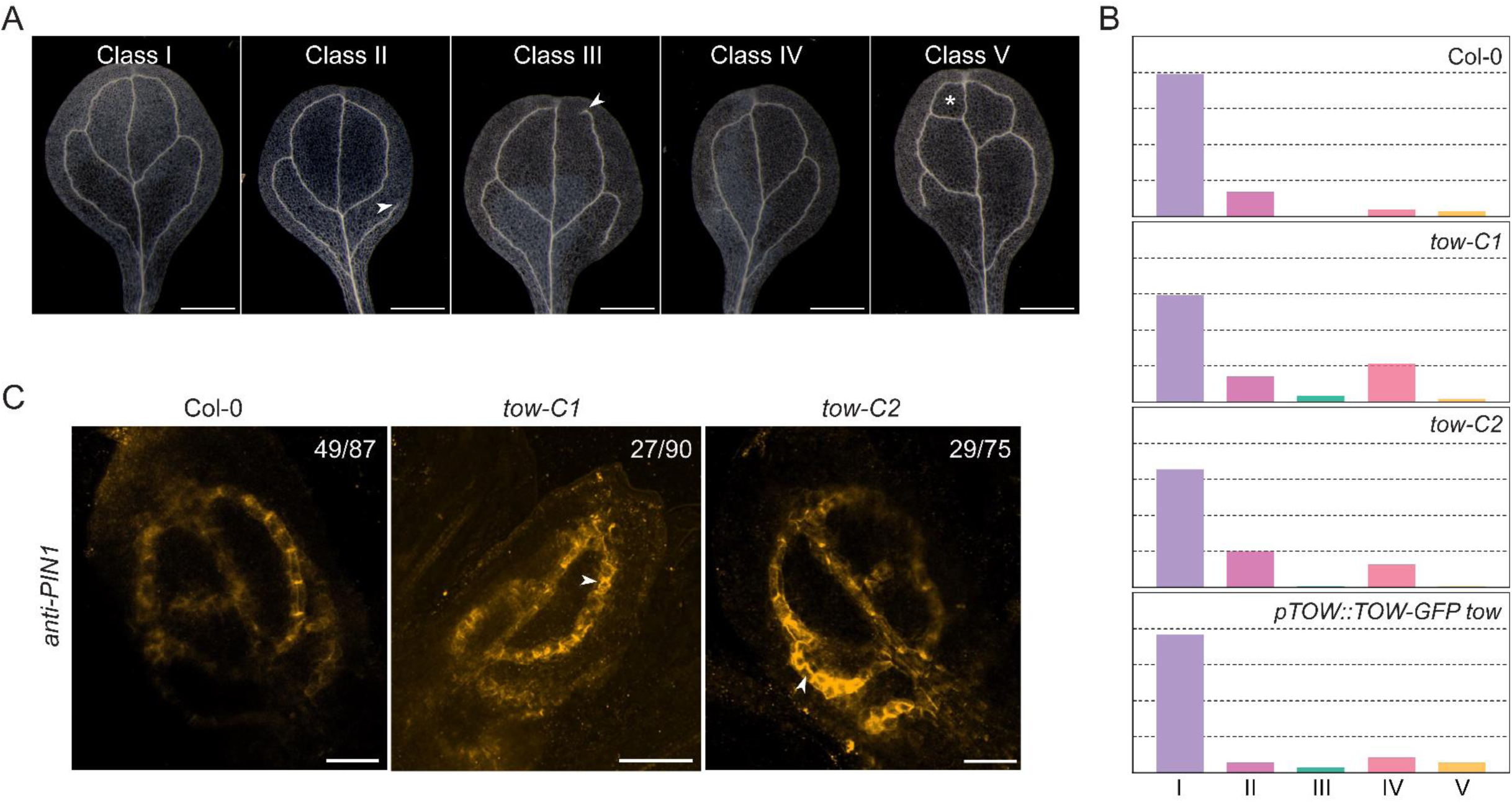
Abnormal patterning and PIN1 polarization during leave venation in *tow* mutants. **(A)** Dark-field illumination of cleared cotyledons illustrating different venation patterning classes: normal venation pattern (Class I), disconnected bottom loop (Class II), disconnected upper loop (Class III), fewer loops (Class IV), and extra loops (Class V). Scale bar: 1 mm. **(B)** Percentage distribution of each venation patterning class in Col-0, *tow* mutants, and the complementation line. Dashed lines indicate 20%, 40%, 60%, and 80% thresholds. For each genotype, n > 120. **(C)** Representative images showing PIN1 immunolocalization in the first leaves. The numbers in the top-left corners represent the incidence of observed PIN1 polarity localization relative to the number of leaves analyzed. White arrowheads indicate PIN1 polarity defects. Scale bar: 100 µm.

Polar auxin transport, mediated by PIN proteins, is the key prerequisite of the canalization mechanism (Sauer et al., 2006). Among the PIN family, PIN1 is the most informative marker for vein patterning, with its expression known to precede and predict vascular strand formation in developing leaves (Scarpella et al., 2006). To determine whether the venation defects in *tow* mutants are associated with altered PIN1 localization, we performed immunolocalization in the first true leaves of Col-0 and *tow* mutants. In Col-0, PIN1 protein was directionally localized toward the leaf tip in early-stage secondary veins connecting to the midvein (49/87 observed) (Figure 2C). In contrast, *tow-C1* and *tow-C2* showed disrupted PIN1 polarity, with significantly fewer cells exhibiting tip-ward localization (27/90 in *tow-C1*, 29/75 in *tow-C2*). Instead, PIN1 signals were often misoriented toward lateral sides - either inward toward the midvein or outward toward the leaf margin (Figure 2C). This loss of coherent PIN1 polarity likely impairs auxin flux and compromises the establishment of continuous vascular strands.

Together, these findings identify a role for TOW in mediating the PIN1 polarization in forming vasculature during leaf vein formation.

### TOW Mediates Vascular Regeneration and Auxin-Induced Vasculature Formation

Next, we assessed vascular regeneration in *tow* mutants. When a transverse incision was introduced into the inflorescence stem, 90% Col-0 plants (n=10) were able to regenerate new vascular strands around the wound site, as visualized by the toluidine blue (TBO) staining (Figure 3A and 3B). In contrast, complete vascular reconnection occurred in only 10% of *tow-C1* (n=10) and 20% of *tow-C2* (n=10) mutants. Notably, 50% of *tow-C1* and 40% of *tow-C2* failed to regenerate vasculature entirely, while the remaining 40% in both lines formed only partial new vessel cells (Figure 3B).

**Figure 3.**
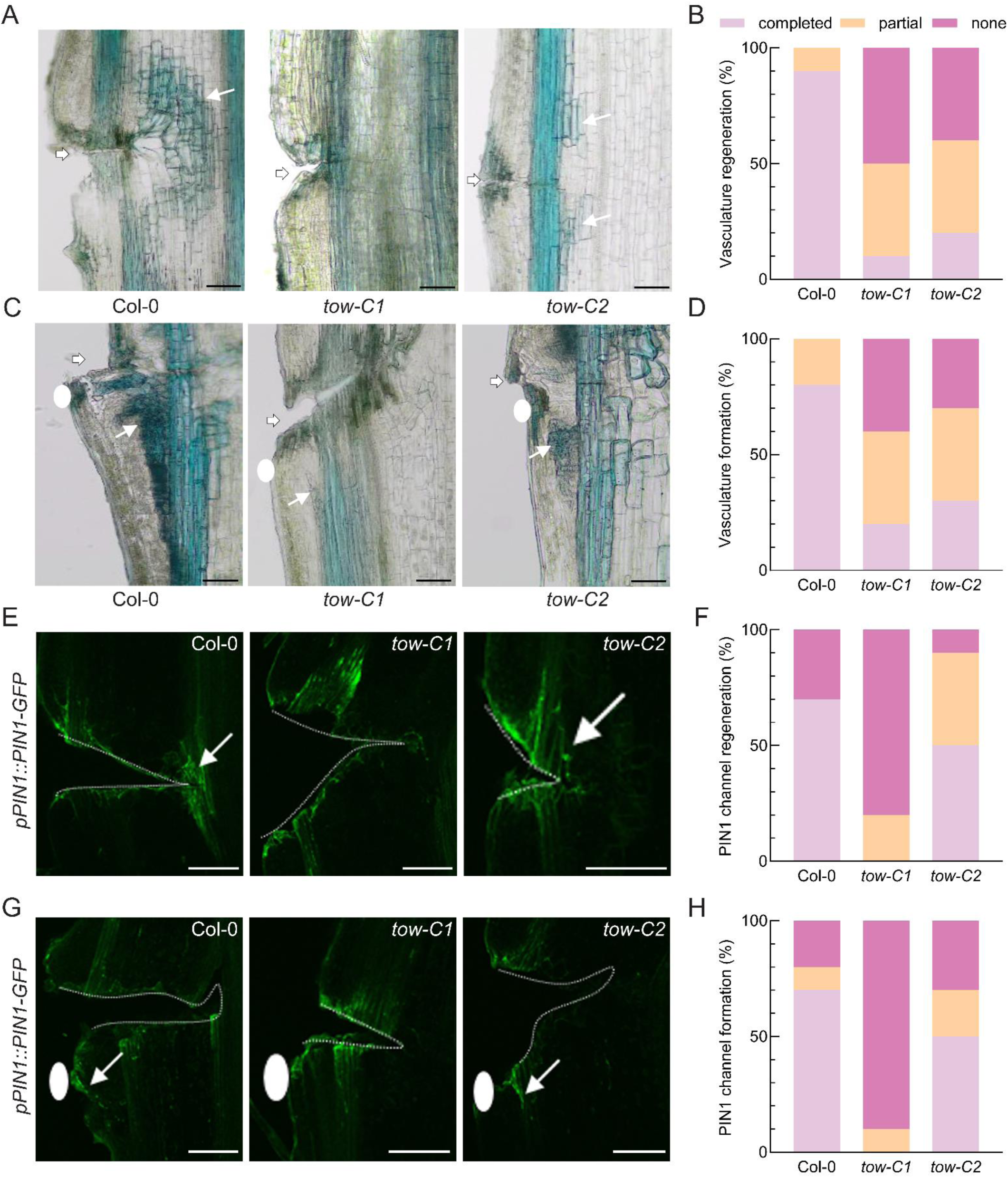
Defective vascular regeneration and auxin canalization in *tow* mutants. **(A-D)** Vascular regeneration (A) or auxin-induced vasculature formation (C) in wounded stems of Col-0 and *tow* mutants with its quantification (B, D). Vasculature is visualized by toluidine blue staining. For each genotype *n* = 10. The wound site is indicated by a white arrowhead, and newly regenerated vessel elements are marked with white arrows. The site of auxin application is marked by a white circle. **(E-H)** Visualization of newly formed PIN1-GFP channels following stem wounding (E) or after local auxin application (G) in Col-0 and *tow* mutants with the quantification (F,H). *n* = 10 for each genotype. The wounding site is outlined with a dashed line, regenerated PIN1-GFP channels are indicated by white arrows. The auxin application site is marked by a white circle. Scale bar: 100 μm

To directly assess auxin canalization potential, we performed an auxin-induced vascular formation assay by excising the apical portion of the stem to remove endogenous auxin sources and applying an IAA droplet just below the wound (Balla et al., 2016; Mazur et al., 2020). In Col-0 (n=10), new vasculature formed in 80% of treated stems, connecting the IAA application site with existing vascular bundles (Figure 3C). In contrast, *tow-C1* (n=10) and *tow-C2* (n=10) showed strong defects in this process, with only 20% and 30% forming continuous vascular strands, respectively. Approximately 40% of *tow-C1* and 30% of *tow-C2* showed completely abolished new vasculature formation, while the remaining 40% in both lines developed only partial connections (Figure 3D).To explore whether these defects correlate with disrupted formation of auxin transport channels, we used the *pPIN1::PIN1-GFP* reporter line. In Col-0 (n=10), polarized PIN1 expression formed a clear channel around the wound in about 70% of samples (Figure 3E and 3F). In *tow-C1* (n=10), no completed PIN1 channel formation was detected; 80% of samples lacked any discernible channel, and 20% showed only partial formation. *tow-C2* mutants (n=10) displayed intermediate defects, with 10% failing to form a channel and 40% forming partial channels (Figure 3F).

Notably, TOW expression was strongly induced upon wounding and local auxin application (Supplemental Figure 4), mirroring PIN1 expression patterns. These findings support a role for TOW in regulating formation of directional PIN1-expressing channels during auxin canalization.

Together, our data demonstrate that TOW is essential for all characterized auxin canalization-associated processes, likely by facilitating the formation of polarized, PIN1-expressing channels required for subsequent vascular patterning.

### TOW Regulates PIN Polarity and Endocytic Trafficking

To investigate whether TOW regulates PIN1 polarity, we first examined auxin-induced PIN repolarization in root, a well-established hallmark of auxin canalization (Sauer et al., 2006). In Col-0 roots, treatment with synthetic auxin NAA promotes the lateralization of basally localized PIN1 to the inner lateral sides of endodermal and pericycle cells (Sauer et al., 2006). This repolarization response was significantly impaired in both *tow-C1* (n=310 cells) and *tow-C2* (n=305 cells) mutants (Figure 4A, 4B). Similar defects were observed for PIN2 lateralization in cortex and epidermal cells (Supplemental Figure 6A, 6C), indicating a general disruption of auxin-induced PIN repolarization in the absence of TOW. These defects were fully rescued in the *pTOW::TOW-GFP tow* complementary line (Supplemental Figure 5A-B, 6A-B), confirming that TOW is required for auxin regulation of PIN polarity.

**Figure 4.**
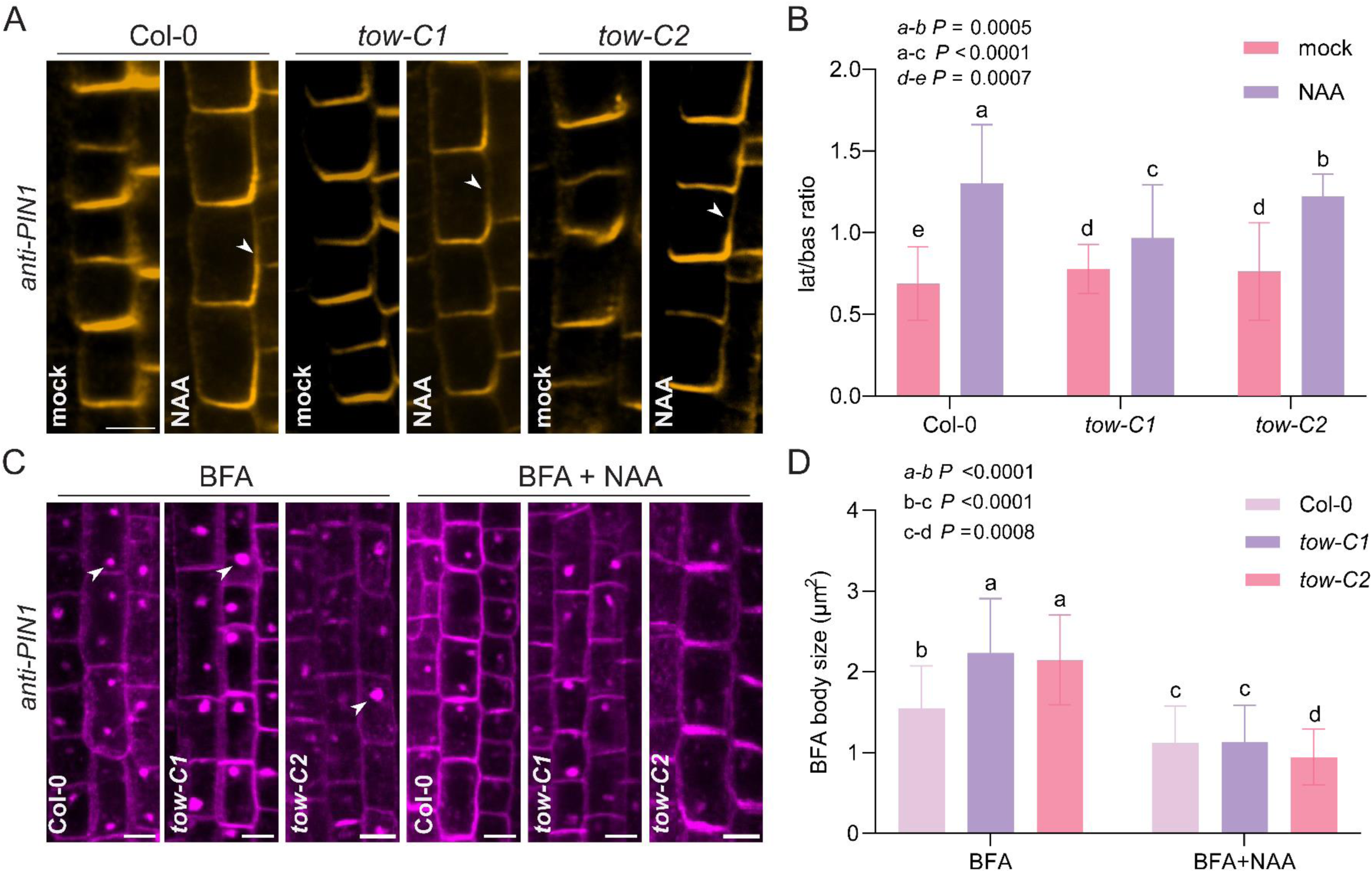
Defective PIN1 polarity and trafficking in *tow* mutants. **(A, B)** Immunolocalization of PIN1 in endodermal cells of the root meristem following treatment with 10 μM naphthaleneacetic acid (NAA) for 4 hr. Arrowheads indicate lateral PIN1 signals. Scale bar, 10 μm. Quantification of lateral-to-basal PIN1 signal intensity (B). **(C, D)** Representative confocal images of BFA bodies in root meristem cells after PIN1 immunostaining. Seedlings were treated with 50 μM brefeldin A (BFA) for 1 hr, or co-treated with 50 μM BFA and 10 μM NAA for 1 hr. PIN1-labled BFA bodies are indicated with arrowheads. Scale bar, 10 μm. Quantification of BFA body size (D) under BFA or BFA + NAA treatment. For (B) and (D), data are presented as mean ± SD from three independent experiments, one representative experiment is shown. Statistical significance was determined using two-way ANOVA followed by Tukey’s post hoc test. Different letters indicate statistically significant differences, *P* < 0.05. n>150 cells for each genotype.

Next, we examined whether TOW influences PIN trafficking. PIN proteins undergo constitutive endocytic recycling, and this can be monitored by treatment with Brefeldin A (BFA), a fungal toxin that inhibits ARF-GEF–mediated vesicle budding. BFA treatment leads to the accumulation of PIN proteins into intracellular aggregates called BFA bodies (Geldner et al., 2001). In contrast, auxin inhibits PIN endocytosis and therefore reduces BFA body formation (Paciorek et al., 2005). After treatment with 50 μM BFA for 1hr, PIN1- and PIN2-labeled BFA bodies were observed in Col-0 (n=236 cells), *tow* mutants, and the complementary line. However, *tow-C1* (n=230 cells) and *tow-C2* (n=230 cells) mutants developed significantly larger BFA bodies than Col-0 (Figure 4C and 4D; Supplemental Figure 6B and 6D), suggesting a defect in PIN trafficking. This phenotype was fully rescued in the *pTOW::TOW-GFP tow* complementary line (Supplemental Figure 5B and 5C), confirming the involvement of TOW in BFA-sensitive PIN trafficking.

**Figure 5.**
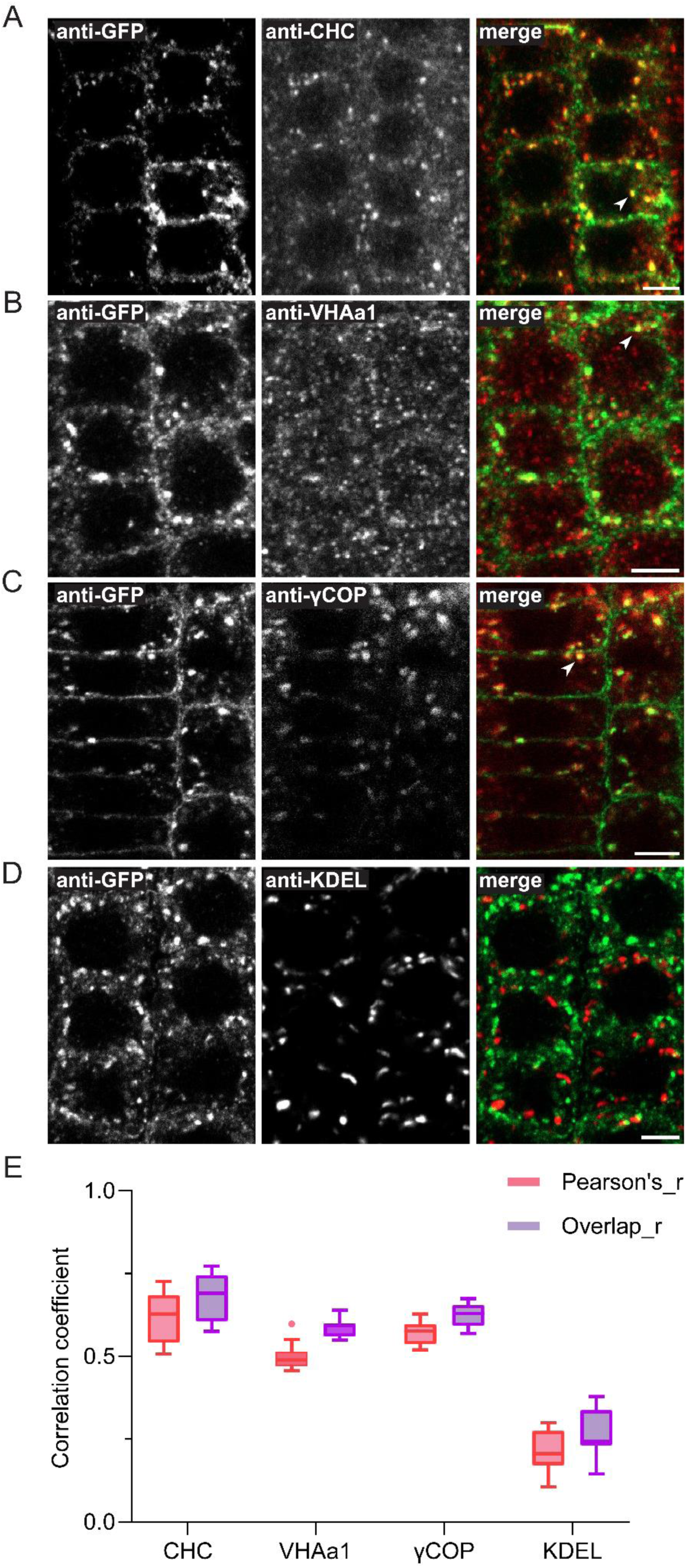
Subcellular Localization of TOW at the Plasma Membrane, TGN, and Golgi. **(A–D)** Immunolocalization of GFP-tagged TOW in the *tow* complementing line co-immunostained with subcellular markers: clathrin heavy chain (CHC) for the trans-Golgi network (TGN) **(A)**, VHAa1 for TGN/early endosomes (TGN/EE) **(B)**, γCOP for the Golgi apparatus **(C)**, and KDEL-containing proteins for the endoplasmic reticulum (ER) **(D)**. For each co-localization experiment, n > 12 roots. Scale bars, 5 μm. **(E)** Quantification of co-localization between *pTOW::TOW-GFP* and each marker. Pearson’s correlation coefficient (Pearson’s_r) and the overlap coefficient (Overlap_r) were calculated from at least 12 roots per marker. Data are presented as a Tukey box plot, where the box represents the interquartile range (25^th^ to 75^th^ percentile), the line inside the box indicates the median, the whiskers extend to the extreme data points within 1.5 times the interquartile range. The individual dot represent outliers beyond this range.

**Figure 6.**
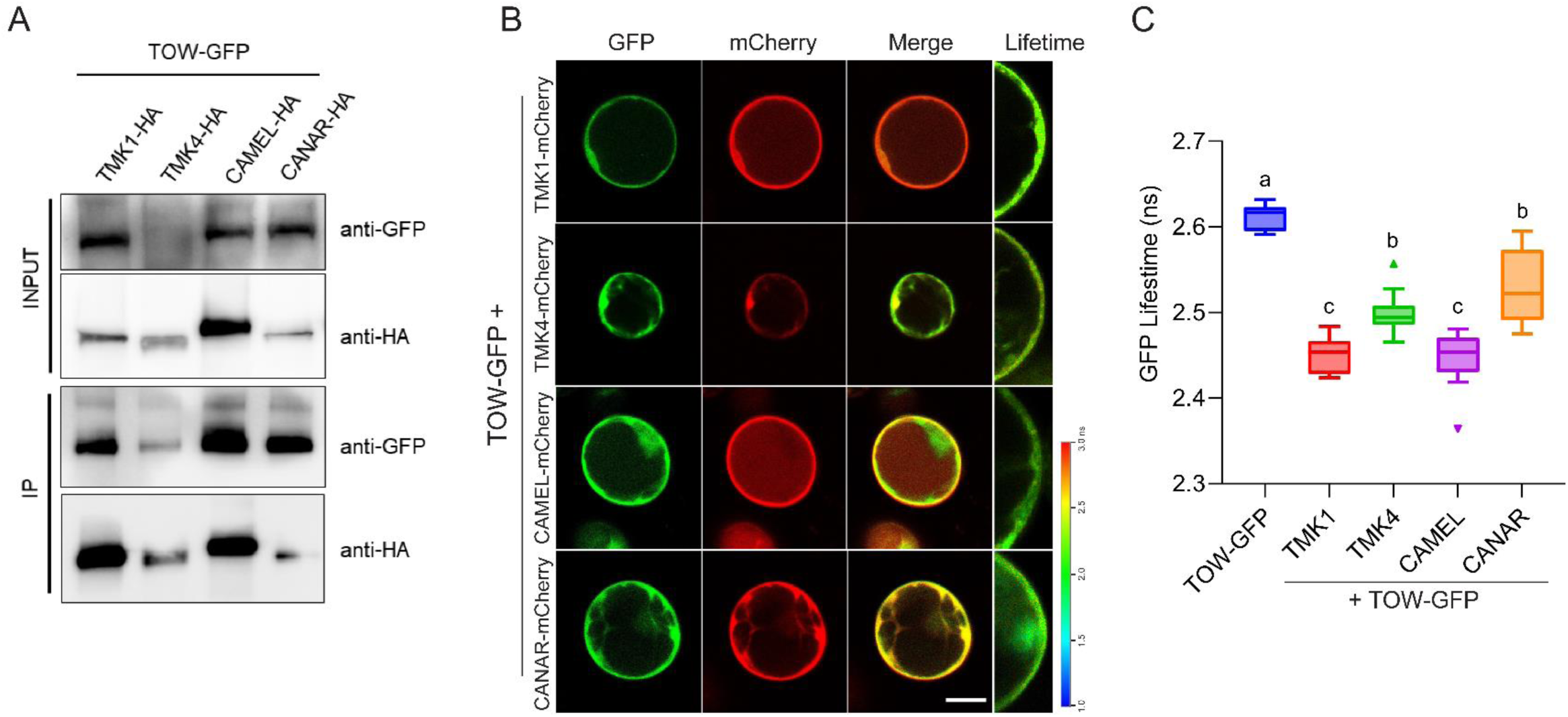
Physical interaction between TOW and RLKs involved in auxin canalization. **(A)** Co-immunoprecipitation (Co-IP) assay showing the interaction between TOW and RLKs (TMK1, TMK4, CAMEL, and CANAR) in Arabidopsis root-derived protoplasts. GFP-tagged TOW and HA-tagged RLKs were transiently co-expressed. Total protein extracts were immunoblotted with anti-GFP and anti-HA antibodies to assess input levels. Co-IP was performed using anti-GFP antibody, and co-precipitated HA-tagged RLKs were detected by immunoblotting with anti-HA. The image shown is representative of two biological replicates. **(B, C)** FRET-FLIM analysis of transiently expressed *35S::TOW-GFP* and *35S::RLKs-mCherry* in protoplasts. GFP fluorescence lifetime was calculated from at least 10 manually selected regions of interest across 12 individual protoplast cells. Heatmaps show fluorescence lifetime values. Data are presented as Tukey box plots from three independent experiments; one representative experiment is shown. Statistical significance was determined using one-way ANOVA followed by Tukey’s post hoc test.

Importantly, the excessive accumulation of PIN proteins in BFA body was significantly reduced by exogenous NAA treatment or BFA washout in both Col-0 and *tow* mutants (Figure 4C and 4D; Supplemental Figure 5E and 5F), further highlighting that TOW is essential for integrating these antagonistic signals to maintain balanced PIN trafficking dynamics.

Together, these findings suggest that TOW is required for both auxin-induced PIN repolarization and for maintaining the PIN endocytic trafficking. The defects in *tow* mutants point to a role for TOW in a regulatory network that controls PIN polarity, potentially by modulating BFA-sensitive trafficking components or interacting with PIN polarity regulators such as kinases.

### Subcellular Localization of TOW at the Golgi, TGN and Plasma Membrane

As previously mentioned, TOW predominantly localizes to the plasma membrane (Figure 1I). However, a portion of TOW was also detected in endomembranes within the cell interior. To further characterize the intracellular compartments where TOW resides, we performed immunolocalization to assess co-localization between TOW-GFP and various organelle markers.

TOW-GFP showed strong co-localization with Clathrin Heavy Chain (CHC; a TGN marker) (Figure 5A), VHAa1 (a TGN and early endosome marker) (Figure 5B), and γCOP (a Golgi marker) (Figure 5C), but not with KDEL-containing proteins (an ER marker) (Figure 5D). Quantitative analysis using Pearson correlation and overlap coefficient confirmed significant overlap between TOW-GFP and CHC, VHAa1, and γCOP, but not with KDEL (Figure 5E). These findings demonstrate that TOW localizes to the plasma membrane, trans-Golgi network (TGN), and Golgi, aligning with key steps in PIN trafficking routes and reinforce TOW involvement in regulation of PIN trafficking.

### TOW Physically Interacts with RLKs Involved in Auxin Canalization

Several RLKs, including CAMEL, CANAR, and TMK1 are well-established components of PIN1-dependent auxin canalization (Hajný et al., 2022). CAMEL, a direct target of WRKY23, functions together with its interactor CANAR as a receptor complex that modulates PIN polarization and auxin flow (Hajný et al., 2020). Similarly, TMK1, a co-receptor of the auxin receptor ABP1, plays an essential role in auxin-guided vascular development (Friml et al., 2022). These RLKs most likely regulate PIN polarity through phosphorylation. Notably, TOW shares overlapping subcellular localization and mutant phenotypes with these RLKs, suggesting a potential functional relationship.

To examine whether TOW physically associates with these RLKs, we co-expressed GFP-tagged TOW and HA-tagged TMK1, CAMEL, or CANAR in Arabidopsis protoplasts. Co-immunoprecipitation using anti-GFP beads revealed that TOW is able to interact with all three RLKs (Figure 6A).

To further validate and quantify these interactions, we performed Förster resonance energy transfer coupled with fluorescence lifetime imaging microscopy (FRET-FLIM). Co-expression of *35S::TOW-GFP* with mCherry-tagged CAMEL, CANAR and TMK1 resulted in a significant reduction of GFP fluorescence lifetime compared to TOW-GFP alone (Figure 6C), indicative of physical interaction. Spatial resolution from FRET-FLIM confirmed that these interactions occur at the cell surface (Figure 6B).

These results demonstrate that TOW physically interacts with key RLK components, namely the CAMEL-CANA complex and the TMK1 component of cell surface auxin perception; all involved in auxin canalization, suggesting that TOW may act within the same signaling modules to regulate auxin-mediated PIN polarity in auxin canalization processes.

## Discussion

Spontaneous and self-organized formation of directional auxin transport channels provides a positional information for flexible formation of vasculature. The underlying canalization hypothesis proposes positive feedback between auxin and PIN auxin transporters as a key prerequisite. In this study, we identify TOW, a previously uncharacterized membrane protein as a novel component of the auxin canalization machinery. Specifically, TOW functions as an essential component of auxin channel formation in the tissues and at the levels of individual cells, it is required for PIN polarity and endocytic trafficking,

TOW has been identified as auxin-inducible gene being transcriptionally regulated downstream of TIR1/AFB-Aux/IAA-ARF canonical auxin signaling and WRKY23 transcription factor. We show that indeed both, ARFs and WRKY23 regulate TOW promotor activity (Figure 1C). Accordingly, *TOW* is expressed in developing vasculature (Figure 1F–H) associated with increased auxin levels and transport and resembling *WRKY23::GUS* expression pattern (Prát et al., 2018). This spatial overlap reinforces the role of TOW in auxin-mediated developmental processes as a part of a conserved auxin canalization module downstream of WRKY23.

Functionally, TOW is required for auxin canalization-related processes including regeneration and *de novo* formation of vasculature from the local auxin source, also for formation of auxin channels during these processes (Figure 2 and 3). Defects in these processes in *tow* mutants are comparable to those observed in *tmk* (Friml et al., 2022)*, camel* and *canar* (Hajný et al., 2020) mutants. At the level of individual cells, TOW, again similarly as shown for CAMEL and CANAR (Hajný et al., 2020), is involved in auxin-induced PIN repolarization (Figure 4). These commonalities support the idea that TOW functions within the same auxin canalization mechanism as these RLKs.

Our data further indicate that TOW modulates BFA-sensitive endocytic trafficking of PIN proteins. This was supported by the subcellular localization of TOW in the key compartments for vesicle sorting and recycling (Figure 5) and increased BFA-induced intracellular PIN accumulation in *tow* mutants (Figure 4, Supplemental Figure 6). Interestingly, this trafficking defect manifested only after BFA treatment but not when co-treated with auxin or after washout. In contrast, Aminophospholipid ATPase3 (ALA3) - a phospholipid flippase previously shown to regulate PIN polarity and vesicle formation (Zhang et al., 2020) - exhibits trafficking defects under all conditions, indicating that ALA3 functions as a general trafficking factor, whereas TOW appears to act more selectively, potentially downstream or in parallel to auxin regulation.

*TOW* encodes a small membrane protein of unknown function, therefore its mechanistic role in canalization remains mysterious. Notably, TOW physically associates with CAMEL, CANAR and TMK1; all RLKs known to coordinate PIN phosphorylation and polarity required for canalization (Figure 6). Given that TMK1 and the CAMEL-CANAR complex regulate PIN polarity via phosphorylation (Hajný et al., 2020; Friml et al., 2022), TOW might either functions as scaffold or stabilize the complexes of these RLKs, regulate their functions or act downstream as an effector of their signaling cascade.

TOW’s broader localization in endomembranes and its role in PIN trafficking suggests that TOW might serve as a functional bridge between RLK-mediated signaling at the plasma membrane and PIN endocytic trafficking. While RLKs such as CAMEL and CANAR likely act through phosphorylation to control PIN localization, TOW appears to translate these upstream signals into trafficking events, especially those regulated by auxin. This unique positioning would allow TOW to ensure the spatial and temporal precision of PIN trafficking necessary for effective polarization of auxin transport for canalization.

In summary, TOW emerges as a novel component involved in auxin canalization. It is extensively regulated downstream of the TIR1/AFB–Aux/IAA–WRKY23 intracellular auxin signaling pathway, while also interacting with TMK1, a key component of cell surface auxin signaling, as well as with the established plasma membrane canalization complex CAMEL–CANAR. Although its precise molecular function remains elusive, TOW likely mediates input from canonical, transcriptional signaling and links cell surface auxin perception with PIN endocytic trafficking, thereby coordinating polarized PIN delivery to the plasma membrane - a mechanism proposed for auxin canalization by modelling approaches (Wabnik et al., 2010). Future studies should focus on elucidating the molecular function of TOW, identifying its functional domains, expanding the characterization of its interactome, and uncovering the nature of mutual regulation between TOW and its associated receptor-like kinases.

## Material and Methods

### Plant Material and Growth Conditions

All *Arabidopsis thaliana* mutants and transgenic lines used in this study are in the Col-0 background. The *arf7 arf19* double mutant (Hajný et al., 2020), *35S::WRKY23-GR* (Prát et al., 2018), and *pPIN1::PIN1-GFP* (Benková et al., 2003) have been described previously.

The *tow* mutants were generated using CRISPR-Cas9 technology (Ge et al., 2019). To enhance editing efficiency, we designed two single-guide RNA (*sgRNA*) sequences targeting *TOW* (*AT3G09280*) gene. *Cas9-free* homozygous mutants were identified through hygromycin resistance screening and confirmed by direct sequencing of PCR products from T3-generation offspring. Four independent mutation events were selected for the test.

To generate marker lines in the *tow* mutant background, *pPIN1::PIN1-GFP* seedlings were crossed with *tow* mutants. Homozygous offspring were selected through fluorescence screening and direct sequencing of PCR products from the T3 generation.

For complementary lines, a 2.5 kb genomic fragment containing the promoter and coding regions of *TOW* was amplified from genomic DNA and cloned into the pDONR221 vector. The resulting entry clones were recombined into the binary vector pB7FWG0 or pB7HAWG0 to generate *pTOW::TOW-GFP*. The final constructs were introduced into *A. tumefaciens GV3101* via electroporation, and subsequently transformed into *tow* mutants using the floral dip method.

To generate the *35S::TOW-GFP* transgenic line, the TOW coding sequence (CDS) without a stop codon was cloned into pDONR221 before being recombined into the destination vector pB7FWG2. The final constructs were then transformed into Col-0 using the floral dip method. All primers used for plasmid construction are listed in Supplementary Table 1.

Seeds were surface-sterilized with chlorine gas, sown on AM+ medium containing half-strength Murashige and Skoog (0.5× MS), 1% (w/v) sucrose and 0.8% (w/v) phytoagar (pH 5.9), stratified in the dark at 4 °C for 2 days, and then grown vertically at 21 °C under a long-day photoperiod (16 h light/8 h dark). The light source consisted of Philips GreenPower LED production modules (deep red [660 nm]/far-red [720 nm]/blue [455 nm] combination, Philips) with a photon density (Li et al., 2021) of 140.4 μmol m−2 s−1 ± 3%.

### Plasmid Construction

For in planta expression, entry vectors containing the coding sequences of TMK1, CAMEL, CANAR, and TMK4 (Tan et al., 2020) were recombined into the pB7ChWG2 binary vectors to generate C-terminal mCherry fusions. The CDS region of *ARF7, ARF19* and *WRKY23* were amplified and cloned into the pENTR/D-TOPO vector, respectively. The resulting entry clones were recombined into the binary vector pB7HAWG2 to generate *35S::ARF7-HA*, *35S::ARF19-HA* and *35S::WRKY23-HA*.

To construct the TOW promoter reporter, an entry vector carrying the 2,156 bp upstream fragment of the *TOW* gene was recombined into the dual luciferase reporter vector pGreenII 0800-LUC (Hellens et al., 2005) to generate the *pTOW:DUAL-LUC* plant binary vector. Primers used for plasmid construction are listed in Supplementary Table 1.

### Quantitative Real-Time PCR

RNA extraction, cDNA synthesis, and reverse transcription-quantitative PCR (RT-qPCR) were performed as described previously(Qi et al., 2022). Briefly, five-day-old seedlings were transferred to AM+ liquid medium (mock) or medium containing IAA or DEX. Each treatment was performed in three biological replicates. Seedlings were collected at the indicated time points after treatment, and RNA was extracted using the RNeasy Plant Mini Kit (QIAGEN, 74904). One microgram of total RNA was used for reverse transcription after genomic DNA removal, following the instructions of the RevertAid First Strand cDNA Synthesis Kit (Thermo, K1622). cDNA was diluted 20-fold before RT-qPCR. Samples were pipetted in three technical replicates using an Automated Workstation Biomek i5 (Beckman Coulter). RT-qPCR was performed with a LightCycler 480 (Roche) using Luna Universal qPCR Master Mix (NEB, M3003S). Sequences of gene-specific primers are listed in Supplementary Table 1. Relative gene expression levels were calculated using the *ΔΔCT* method, with *PROTEIN PHOSPHATASE 2A SUBUNIT A3 (PP2AA3)* as the internal control.

### TOW Promoter Activation and Dual Luciferase Assay

For promoter activation assays, the *pTOW:DUAL-LUC* construct was transiently co-expressed in *Nicotiana benthamiana* with free HA or HA-fused *ARF7*, *ARF19*, and *WRKY23*. Specifically, *A. tumefaciens* GV3101 strains carrying expression constructs *35S::WRKY23-HA*, *35S::ARF7-HA*, *35S::ARF19-HA*, empty pB7HAWG2, and *pTOW:DUAL-LUC* were grown overnight in LB medium, pelleted by centrifugation, and resuspended in infiltration buffer (10 mM MES, 10 mM MgCl₂, and 200 µM acetosyringone) to an OD600 of 1.0. Equal volumes of different constructs were mixed for infiltration. Cultures were spot-infiltrated into four-week-old tobacco leaves. Leaf discs were collected two days after infiltration, ground in liquid nitrogen, and lysed with PLB buffer from the Dual-Luciferase Reporter Assay System (Promega, E1910). Lysates were centrifuged at 12,000 g for 1 min, and 10 μL of supernatant was used to measure FLUC and RLUC activities according to the manufacturer’s instructions using a Victor3 plate reader (PerkinElmer). FLUC and RLUC substrates were added via an automatic injector at 25°C, followed by 2 s of shaking with a 2 s delay. Signals were captured for 3 s and recorded as counts per second. To quantify *TOW* promoter activity, the ratio of FLUC to RLUC activity was calculated for each combination.

### *in situ* immunolocalization

Immunostaining in primary root was performed on three-day-old seedlings as previously described (Sauer et al., 2006). The primary antibodies used were rabbit *anti*-PIN1 (Hajný et al., 2020), diluted 1:1,000 (v/v); rabbit *anti*-PIN2 diluted 1:1,000 (v/v) (Hajný et al., 2020), mouse anti-GFP (cat. no. AS20 4511; Agrisera), diluted 1:5,000 (v/v); rabbit anti-CHC (cat. no. A304-743A, ThermoFisher), diluted 1:1,000 (v/v); rabbit *anti*-γCOP (cat. no. AS08 327; Agrisera) diluted 1:1,000 (v/v); rabbit anti-VHA a1, rabbit *anti*-KDEL (cat. no. AS08 325; Agrisera) diluted 1:1,000 (v/v). The secondary antibody used was sheep anti-rabbit conjugated with Cy3 (cat. no. C2306; Sigma-Aldrich), diluted 1:600 (v/v) and anti-mouse Alexa Fluor 488 (cat. no. A11029; Sigma-Aldrich), diluted 1:600 (v/v). For PIN1 immunolocalization in young leaves, tissues were cleared sequentially in methanol and ethanol/xylene solution after fixation, and subsequent procedures were carried out as described for roots.

### Cotyledon Vasculature Analysis

Cotyledons of 10-day-old seedlings were harvested and incubated overnight in 70% ethanol for initial clearing. The samples were then transferred to a 4% (v/v) HCl and 20% (v/v) methanol solution and incubated at 65°C for 15 minutes, followed by incubation in a 7% (v/v) NaOH and 70% (v/v) ethanol solution at room temperature for 15 minutes.

To ensure gradual rehydration, cotyledons were sequentially transferred through an ethanol series of 70% (v/v), 50% (v/v), 25% (v/v), and 10% (v/v) ethanol, with each step lasting 5 minutes. The samples were then immersed in a 25% (v/v) glycerol and 5% (v/v) ethanol solution before being mounted in 50% (v/v) glycerol for imaging. Differential interference contrast (DIC) microscopy was performed using an Olympus BX53 microscope.

### Local Auxin Application and Vascular Regeneration Experiments in Arabidopsis Stems

Young Arabidopsis plants with inflorescence stems at the primary tissue stage (vascular bundles separated by interfascicular parenchyma) were selected for a two-step experiment, as previously described (Mazur et al., 2016). First, the flowering parts of the stems were removed using a sharp razor blade, leaving a 7 cm-long stem. To stabilize the stems, they were attached to a polypropylene tube and subjected to a 2.5 g lead weight for six days, promoting the formation of a closed ring of cambium around the stem circumference(Mazur et al., 2014). Next, a transverse incision was made above the leaf rosette to disrupt the longitudinal cambium continuity and the basipetal transport of endogenous auxin. A droplet of lanolin paste containing IAA (Sigma-Aldrich, cat. no. 15148-2G) was then applied just below the cut. This setup ensured that any observed changes resulted solely from the externally applied auxin. The lanolin-IAA mixture was refreshed every two days throughout the experiment. Each experiment was conducted twice per plant line, with at least 10 plants analyzed per replicate. Finally, the samples were collected, manually sectioned, and mounted in a 50% glycerol aqueous solution for imaging.

### Förster Resonance Energy Transfer with Fluorescence Lifetime Imaging Microscopy (FRET-FLIM)

Protoplasts were isolated from root cell suspension cultures as previously described (Hajný et al., 2020). Plasmids were prepared using the E.Z.N.A. Plasmid Maxi Kit I (Omega Bio-Tek). Protoplasts were transfected with 10 µg of plasmid DNA using the PEG-calcium-mediated transformation method and incubated in the dark at room temperature for 12–16 hr before imaging. FRET-FLIM measurements were performed using a Leica TCS SP8 confocal microscope equipped with a PicoQuant FLIM system. The donor fluorophore was excited at 488 nm using a 70% white-light laser source (10% intensity), and emission fluorescence was collected by a HyD detector with bandpass filters of 499–551 nm and 600–650 nm. The acceptor fluorophore was excited at 561 nm using the same 70% white-light laser source (50% intensity), and emission was collected using a 600–650 nm bandpass filter. Time-correlated single-photon counting (TCSPC) data acquisition was performed using PicoQuant SymPhoTime software, with an acquisition time of 60–120 seconds per image to ensure sufficient photon counts (>1,000 photons per pixel). A segmented line was drawn along the plasma membrane (PM) region to measure the mean signal intensity for each channel. Fluorescence decay curves were analyzed using Leica LAS X software, and lifetime values were extracted from at least 10 independent regions of interest (ROIs) per sample. Statistical significance was assessed using one-way ANOVA followed by Tukey’s post hoc test in GraphPad Prism. To generate the FRET-SE efficiency heatmap, the corresponding ROIs were processed using the Image FLIM module in LAS X.

### Co-Immunoprecipitation Assay

Co-immunoprecipitation assays were performed using Arabidopsis root protoplasts. Transfected protoplasts were lysed in 500 µL of lysis buffer (50 mM Tris-HCl, pH 7.4, 150 mM NaCl, 1 mM DTT, 1× protease inhibitor cocktail (cOmplete Tablets, Roche)). The membrane fraction was isolated by flash-freezing the protoplasts in liquid nitrogen, thawing on ice, and centrifuging at 14,000 *g* at 4°C for 45 minutes. The supernatant was discarded, and the pellet was resuspended in lysis buffer supplemented with 0.5% NP-40. The sample was then centrifuged at 12,000 *g* at 4°C for 15 minutes, and the supernatant was collected for co-immunoprecipitation. For co-immunoprecipitation, the membrane fractions were incubated with anti-GFP beads (Miltenyi Biotec) for 1 hr at 4°C. The immunoprecipitated proteins were washed four times with Washing Buffer 1 (150 mM NaCl, 1% Igepal CA-630, 0.5% sodium deoxycholate, 0.1% SDS, 50 mM Tris-HCl, pH 8.0) and once with Washing Buffer 2 (20 mM Tris-HCl, pH 7.5). Proteins were eluted from the beads using preheated elution buffer (50 mM Tris-HCl, pH 6.8, 50 mM DTT, 1% SDS, 1 mM EDTA, 0.005% bromophenol blue, and 10% glycerol) at 95°C. Eluted proteins were separated by 10% (v/v) SDS-PAGE and detected using an anti-GFP-HRP antibody (1:10,000 dilution, cat. no. A8592, Sigma-Aldrich) or an anti-HA-peroxidase antibody (1:10,000 dilution, cat. no. 12013819001, Sigma-Aldrich).

### Imaging via Confocal Laser Scanning Microscopy

For BFA treatment, seedlings were incubated in liquid AM+ medium supplemented with 50 μM BFA or together 10 μM NAA for 1 hr before washing with fresh AM+ medium for 30min. BFA bodies in whole *z*-stacks were analyzed using ImageJ. For colocalization evaluation, *pTOW::TOW-GFP* was co-immunostained with TGN, Golgi or ER markers. At least 12 individual roots were selected for each combination. Pearson correlation and Overlap coefficients were calculated using the Colocalization Finder plugin in ImageJ.

Fluorescence imaging was performed using a Zeiss LSM800 confocal laser-scanning microscope with the following excitation wavelength parameters: Cy3, 548 nm; Alexa Fluor 488, 488 nm.

### Homology Analysis

The TOW protein sequence and its orthologous sequences were obtained from Phytozome 13 and aligned using ClustalW (Thompson et al., 1994). Neighbor-joining (NJ) phylogenetic analysis was conducted in MEGA 11 (Tamura et al., 2021) using protein Poisson distances and pairwise deletion of gap sites. Reliability of the phylogenetic tree was evaluated by performing 1000 bootstrap replicates.

### Software and Statistical Analysis

Data were analyzed using one-way ANOVA followed by Tukey’s post hoc test or two-way ANOVA and Tukey’s post hoc test in GraphPad Prism 8.

### External Data Sources

Arabidopsis gene and protein sequences are available from TAIR (https://www-arabidopsis-org.libraryproxy.ista.ac.at/). Sequences for *TOW* orthologous genes were obtained through a BLAST search with the AtTOW protein sequence in Phytozome 13 (https://phytozome-next.jgi.doe.gov/blast-search).

## Acknowledgments

We thank Dr. Z. Ge (ISTA) for providing vectors for the *CRISPR/Cas9* system, Prof. Michael Wrzaczek (Czech Academy of Sciences, Czechia) for valuable suggestions and Prof. Maciek Adamowski (University of Gdańsk) for technical assistance. We also acknowledge the support of the Imaging & Optics Facility and the Lab Support Facility at the Institute of Science and Technology Austria. The authors declare no conflict of interest.

## Author Contribution

Conceptualization, M. Li., and J. F.; Methodology, M. Li., and J. F.; Investigation, M. Li., N.R., G.M., and E.M; Writing – Original Draft, M. Li., and J. F; Writing – Review & Editing, M. Li., and J. F; Funding Acquisition and Resources, M. Li., and J. F.; Supervision, M. Li., and J. F.

---

**Supplemental Figure 1.**
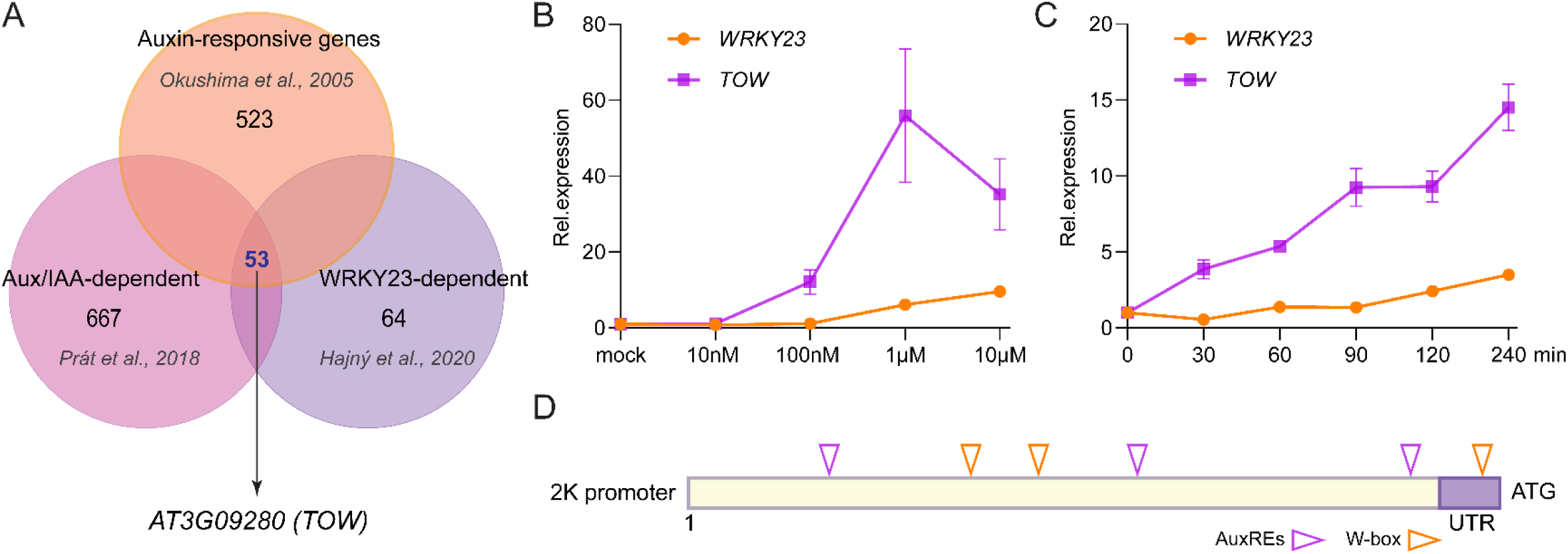
TOW is expressed downstream of SCF^TIR1^-Aux/IAA-WRKY23 signaling pathway. **(A)** Schematic representation of the data mining strategy based on the microarray dataset. **(B, C)** Quantitative RT-PCR analysis showing auxin-dose-dependent upregulation of *TOW* and *WRKY23* transcripts after 4 hr of treatment. (B) and time-dependent upregulation of *TOW* and *WRKY23* expression in response to 100 nM IAA (C). Gene expression levels were normalized to *PP2A*. Data are presented as mean ± SD from three biological replicates; one representative experiment is shown. **(D)** Representation of the 2 kb *TOW* promoter, with triangle markers indicating the locations of auxin-responsive elements (AuxREs) and WRKY-binding motifs (W-box).

**Supplemental Figure 2.**
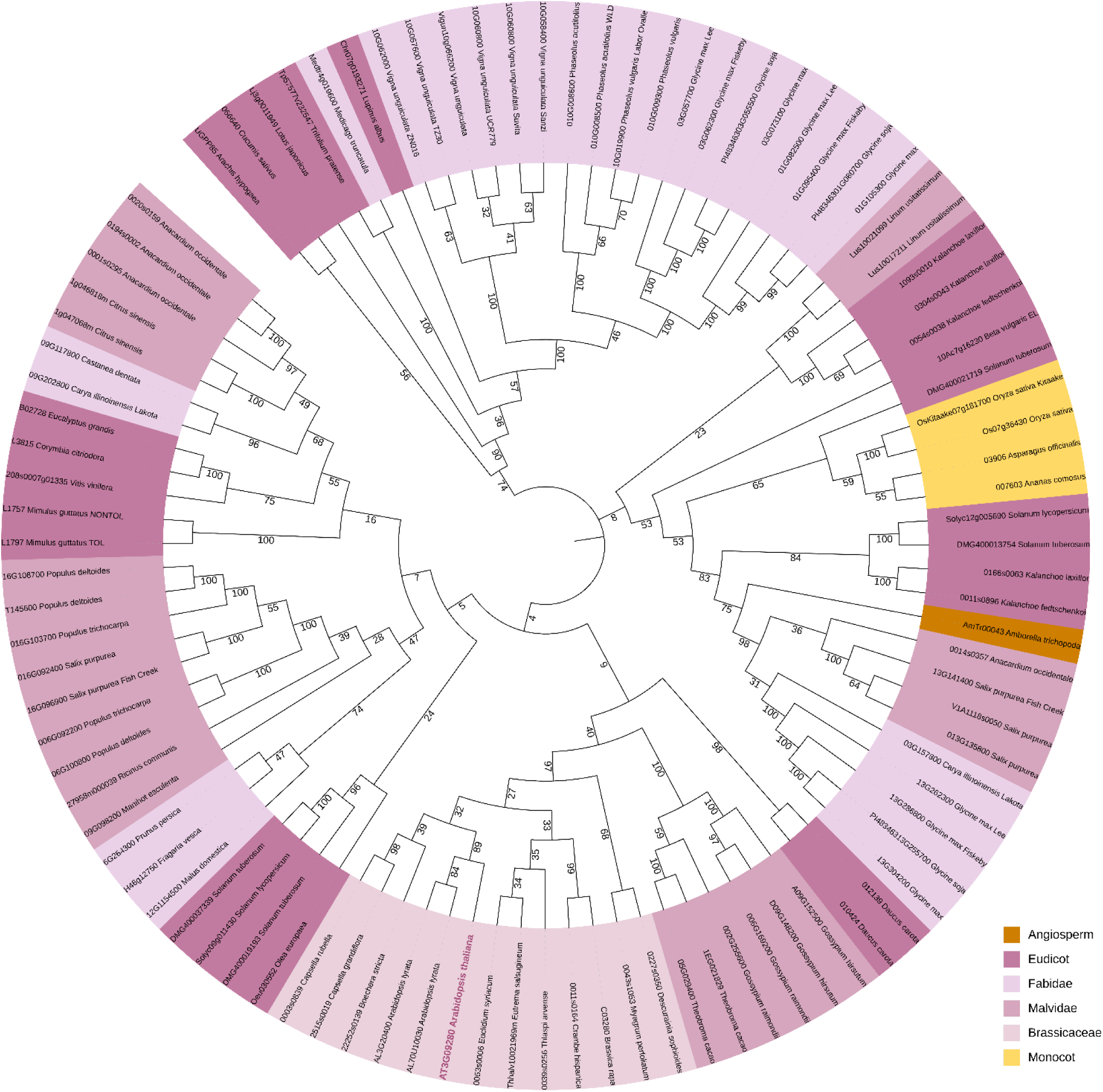
Phylogenetic tree based on the TOW protein sequence. Amino acid sequences of TOW orthologs from various plant groups (algae, mosses, ferns, and seed plants) were retrieved from the Phytozome database. Sequence alignment was performed using ClustalW with the MUSCLE method, and phylogenetic analysis was conducted using MEGA11 with the Neighbor-Joining method, including 1000 bootstrap replicates. The generated phylogenetic tree was visualized using iTOL, with plant species annotations represented in color.

**Supplemental Figure 3.**
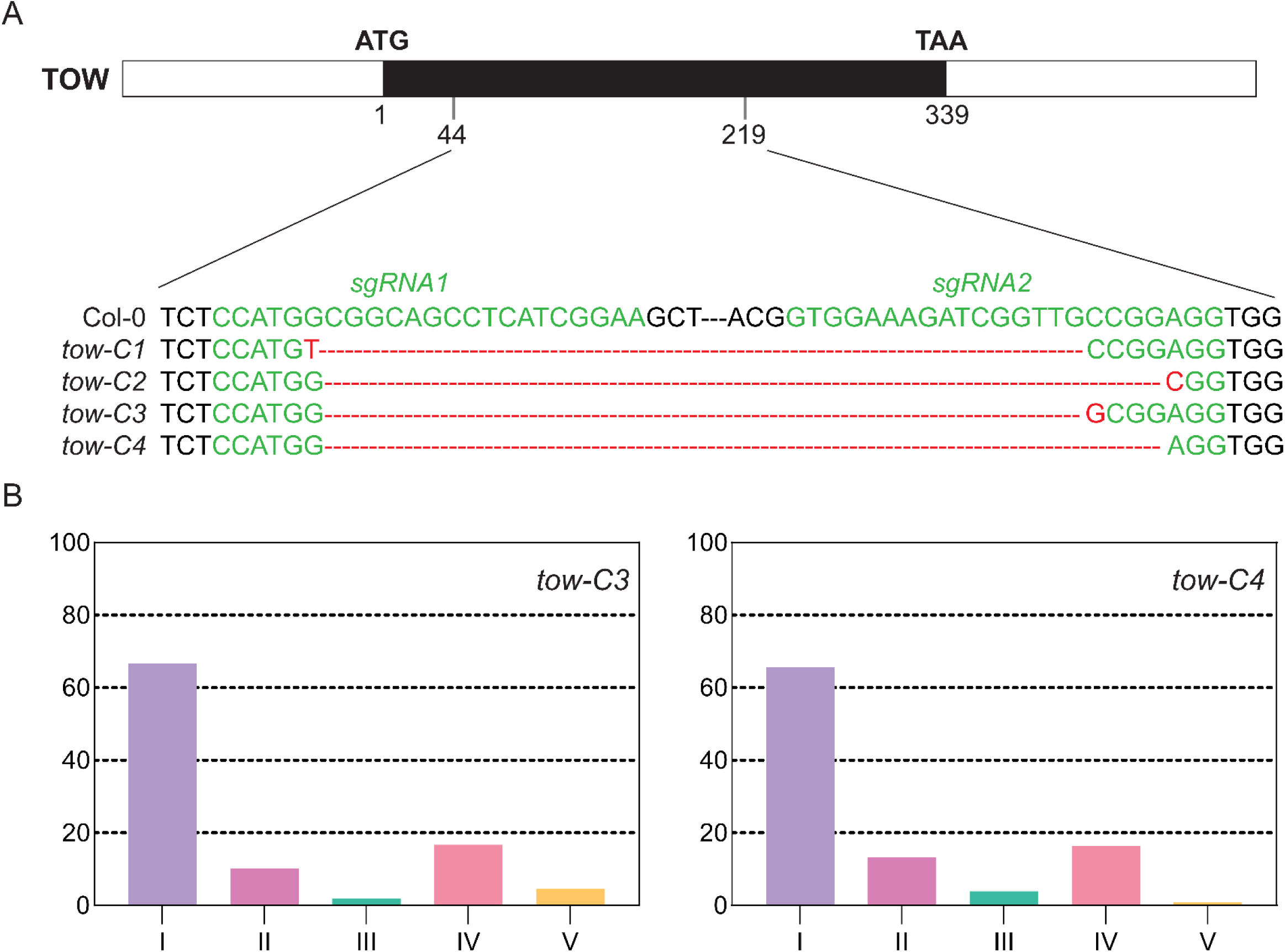
Generation and characterization of *tow* mutants. **(A)** Schematic representation of CRISPR/Cas9 target sites in the *TOW* gene and sequence alignments of *tow* mutant alleles. Black boxes indicate coding exons, and white boxes represent untranslated regions (UTRs). *tow-C1* to *tow-C4* are four independent mutant lines. *tow-C1* and *tow-C3* carry in-frame deletions, while *tow-C2* and *tow-C4* harbor frameshift mutations that result in premature stop codons. **(B)** Quantification of venation patterning phenotypes in *tow-C3* and *tow-C4*. Venation classes include: Class I (normal pattern), Class II (disconnected bottom loop), Class III (disconnected upper loop), Class IV (fewer loops), and Class V (extra loops). *n* > 120 cotyledons per genotype.

**Supplemental Figure 4.**
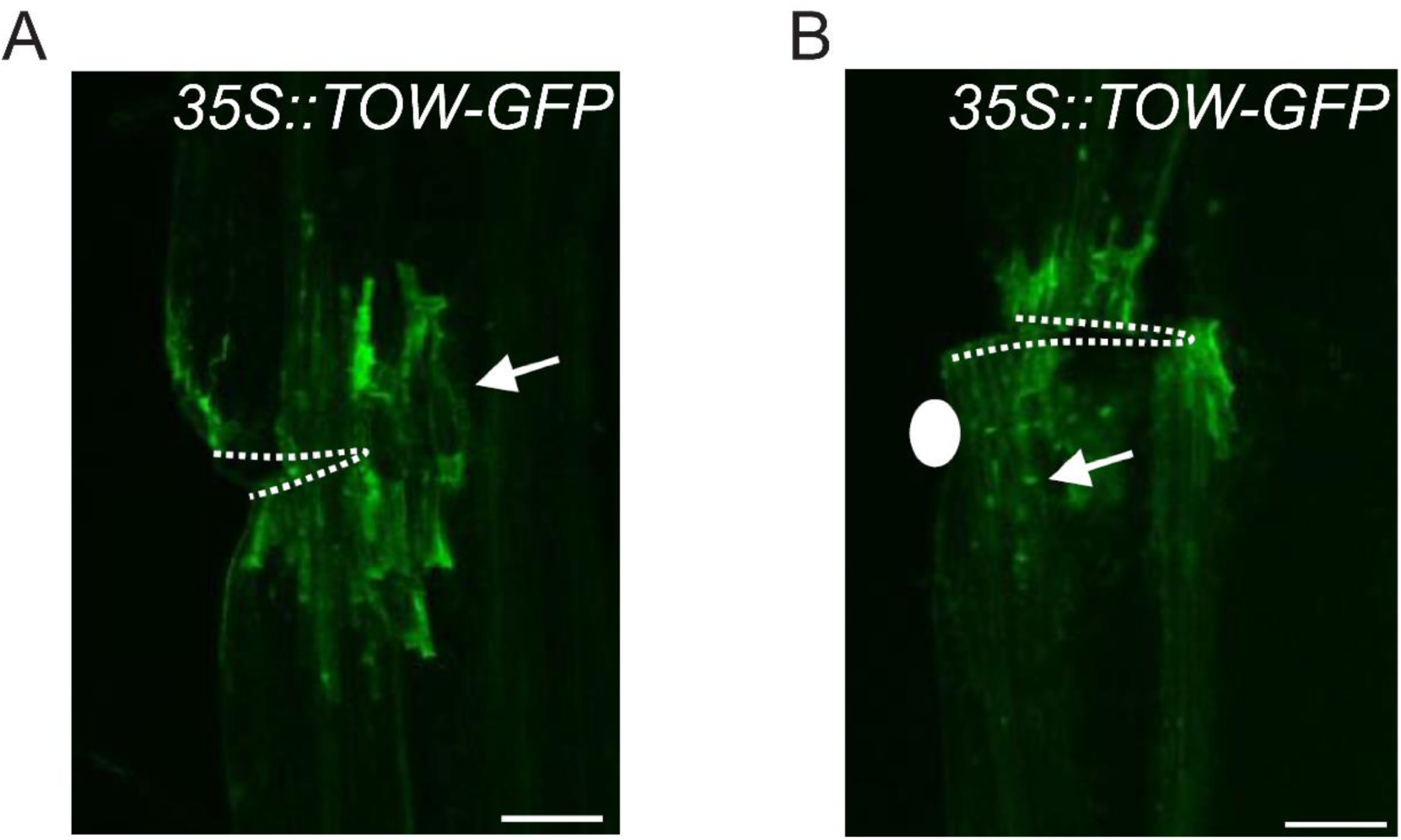
Expression of *35S::TOW-GFP* in stems after wounding **(A)** and local auxin application **(B).** The wounding site is outlined with a dashed line, and the auxin application site is marked by a white circle. TOW-GFP expression is indicated by white arrows. *n* = 10 stems per experiment. Scale bar, 100 μm.

**Supplemental Figure 5.**
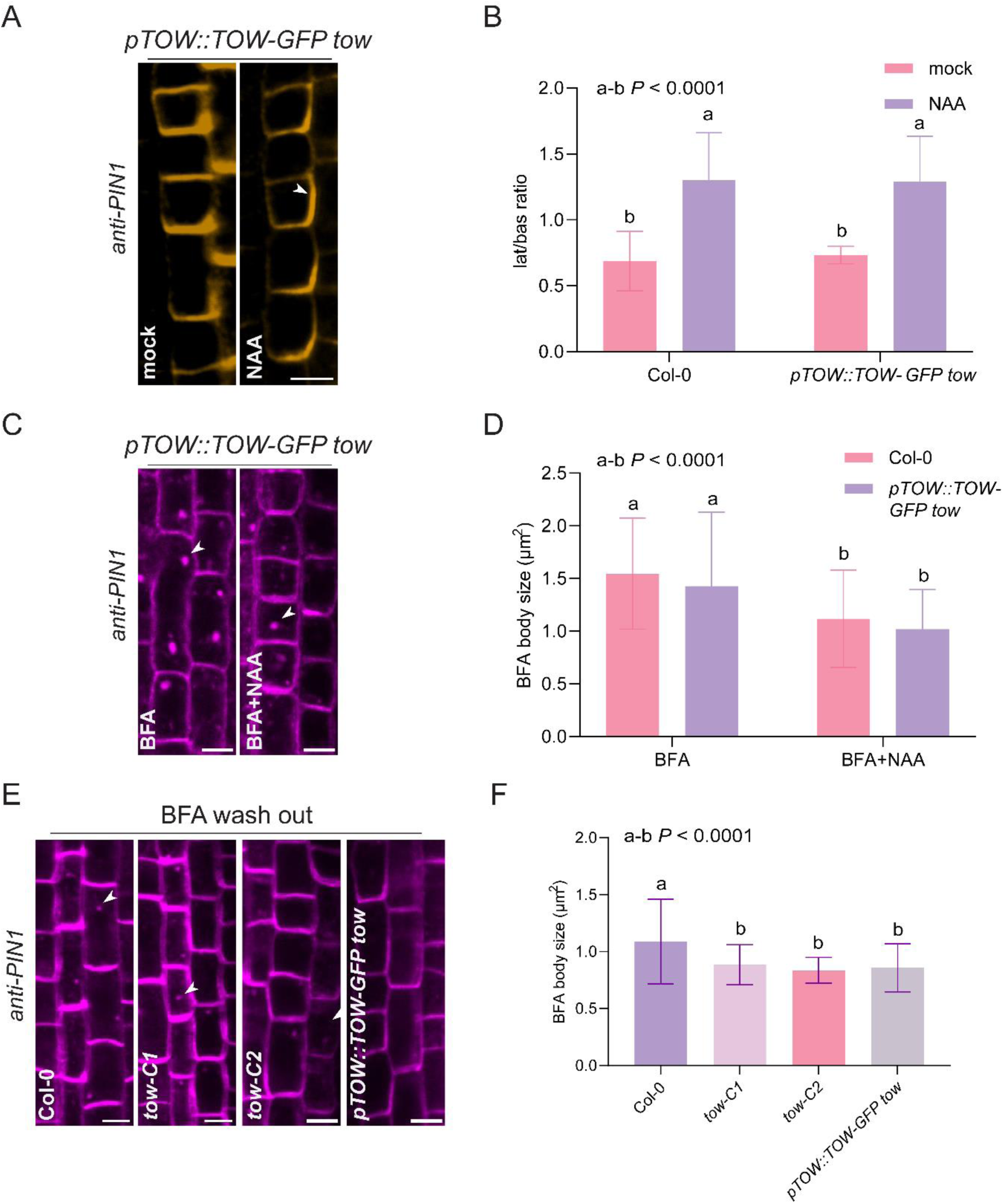
Restoration of PIN1 polarity and trafficking in *pTOW::TOW-GFP tow* line. **(A, B)** PIN1 immunolocalization in root endodermal cells of the *tow* complementary line after treatment with 10 μM naphthaleneacetic acid (NAA) for 4 hr. Arrowheads indicate lateral PIN1 signals. Scale bar, 10 μm. Quantification of lateral-to-basal PIN1 signal ratio (B). **(C, D)** Representative confocal images of PIN1-labeled BFA bodies in the complementary line. Seedlings were treated with 50 μM brefeldin A (BFA) for 1 hr, or co-treated with 50 μM BFA and 10 μM NAA. Arrowheads indicate BFA bodies. Scale bar, 10 μm. Quantification of BFA body size (D). **(E, F)** PIN1-labeled BFA bodies in Col-0, *tow* mutants, and the complementation line after 50 μM BFA treatment followed by washout. Arrowheads mark remaining BFA bodies. Scale bar, 10 μm. Quantification of BFA body size (F). For (B), (D) and (F), data are presented as mean ± SD from three independent experiments, one representative experiment is shown. Statistical significance was determined using two-way ANOVA or one-way ANOVA followed by Tukey’s post hoc test. Different letters indicate statistically significant differences, *P* < 0.05. n>150 cells for each genotype.

**Supplemental Figure 6.**
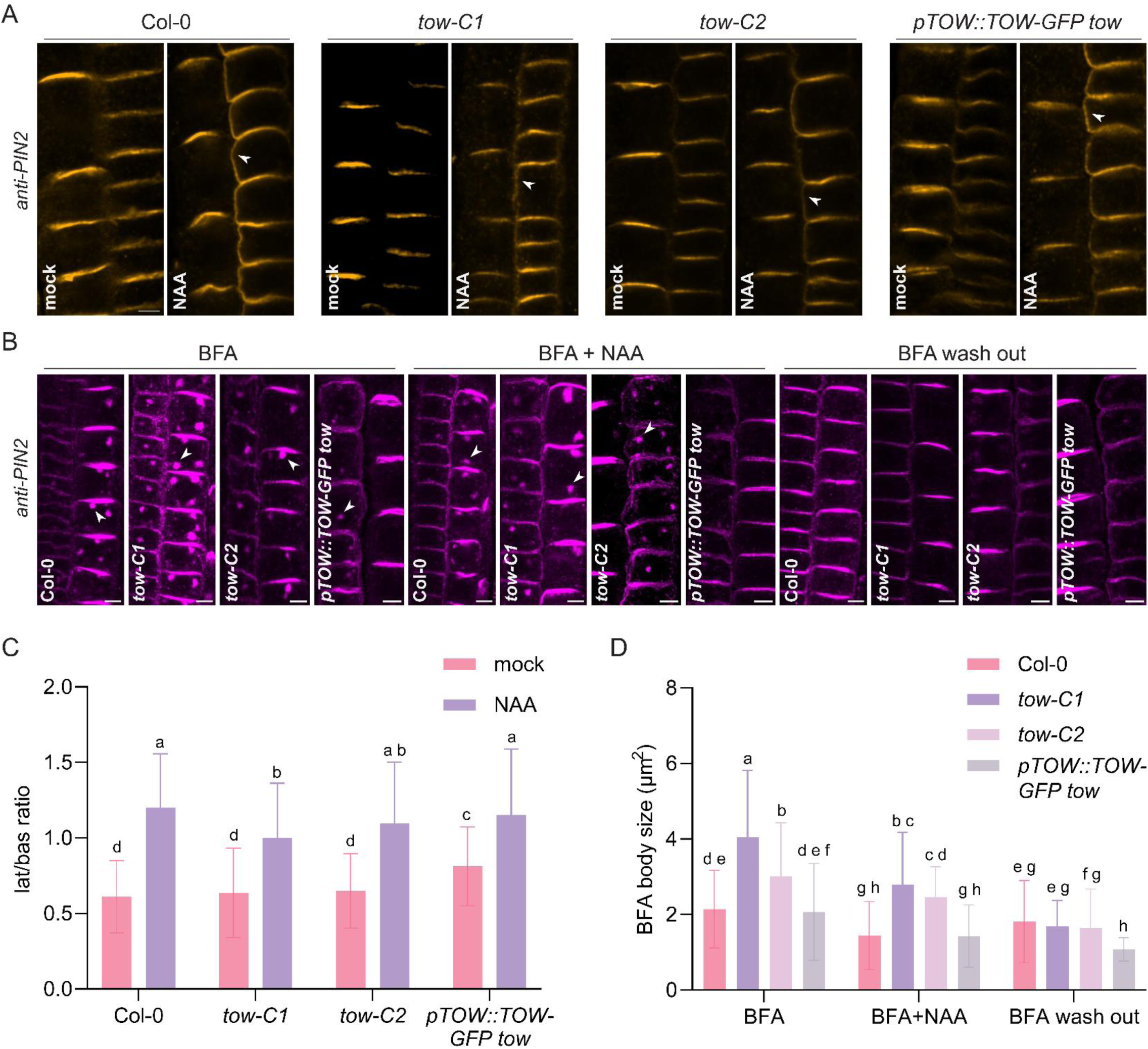
Defective PIN2 polarity and trafficking in *tow* mutants. **(A)** Immunolocalization of PIN2 in cortex and epidermal cells of the root meristem following treatment with 10 μM naphthaleneacetic acid (NAA) for 4 hr. Arrowheads indicate lateral PIN2 signals. Scale bar, 10 μm. **(B)** Quantification of lateral-to-basal PIN2 signal intensity from (A). Data are presented as mean ± SD from three independent experiments; one representative experiment is shown. Statistical significance was determined using two-way ANOVA followed by Tukey’s post hoc test. Different letters denote statistically significant differences, *P* < 0.05. **(C)** Representative confocal images of BFA bodies in cortex and epidermal cells after PIN2 immunostaining. Seedlings were either treated with 50 μM brefeldin A (BFA) for 1 hr, or co-treated with 50 μM BFA and 10 μM NAA for 1 hr, or followed by 30min washout. Scale bar, 10 μm. **(D)** Quantification of BFA body size shown in (C). Data are presented as mean ± SD from three independent experiments; one representative experiment is shown. Statistical significance was determined using two-way ANOVA followed by Tukey’s post hoc test. Different letters indicate statistically significant differences, *P* < 0.05.

**Supplementary Table 1.**
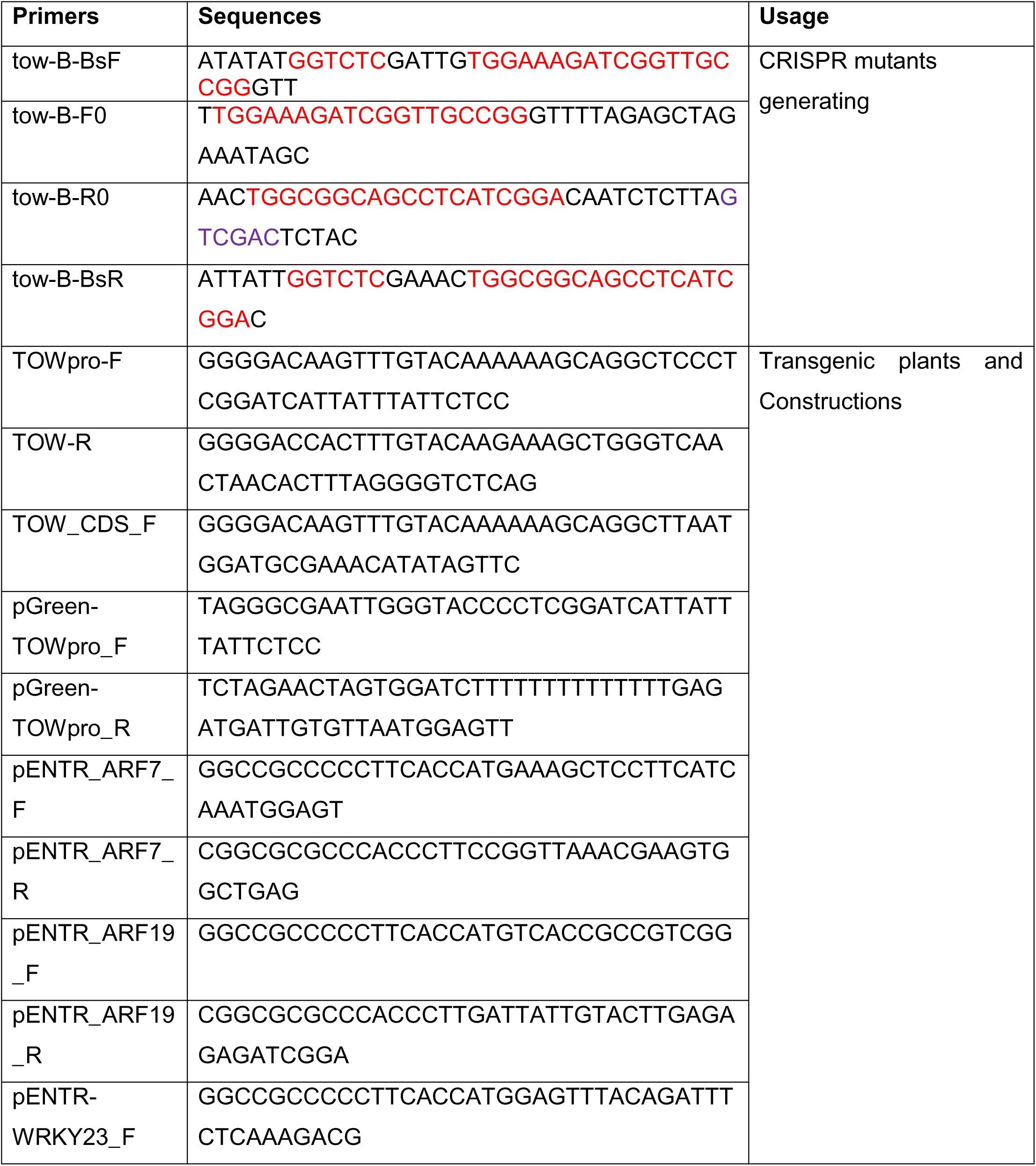
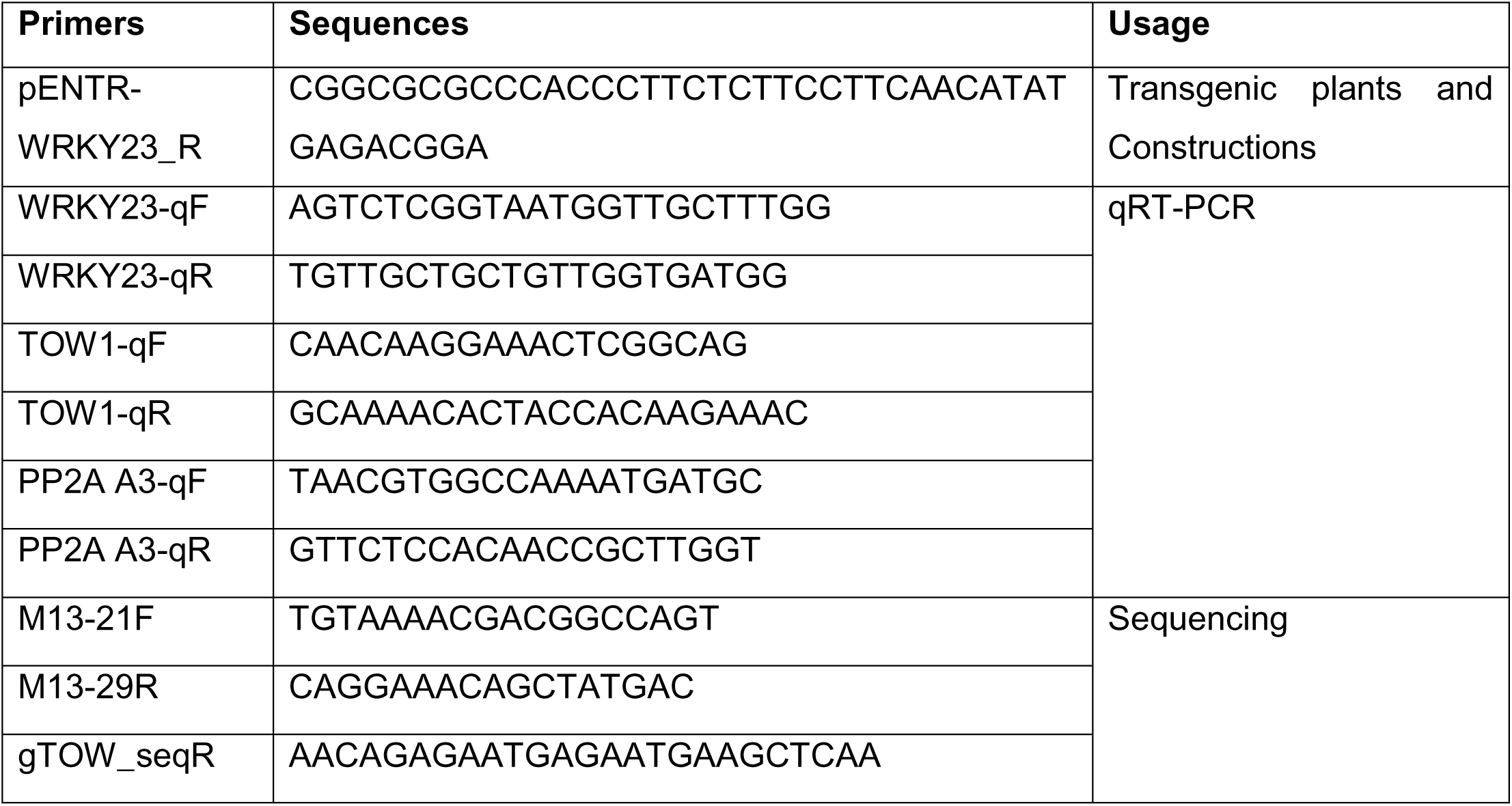
Primers used in this study.

